# Sweet anticipation: Predictability of familiar music in autism

**DOI:** 10.1101/2020.08.03.233668

**Authors:** Patricia Alves Da Mota, Eloise A Stark, Henrique M Fernandes, Christine Ahrends, Joana Cabral, Line Gebauer, Francesca Happé, Peter Vuust, Morten L Kringelbach

## Abstract

Autism has been characterised by different behavioural and cognitive profiles compared to typically developing (TD) individuals, and increasingly these differences have been associated with differences in structural and functional brain connectivity. It is currently unknown as to whether autistic and TD listeners process music in the same way: emotionally, mnemonically, and perceptually. The present study explores the brain’s dynamical landscape linked to music familiarity in an fMRI dataset from autistic and TD individuals. Group analysis using leading eigenvector dynamics analysis (LEiDA) revealed significantly higher probability of occurrence of a brain network in TD compared to autistic individuals during listening to familiar music. This network includes limbic and paralimbic areas (amygdala, hippocampus, parahippocampal gyrus, and temporal pole). No significant differences were found between autistic and TD individuals while listening to a scrambled, i.e. unfamiliar and more unpredictable, version of the same music track. These findings provide novel neuroimaging insights into how autistic prediction monitoring may shape brain networks during listening to familiar musical excerpts.

## Introduction

Music is a universal feature of human societies, represents a physical manifestation of the internal creative impulse in humans, and has been linked to both moment-by-moment pleasure, and overall wellbeing (Eloise A. Stark, Vuust, & Kringelbach, 2018). Among the things that people report as most pleasurable in life, music is consistently among the top ten (Dube & Lebel, 2003), and the ability to derive pleasure from music appears to be a unique trait among the human species (Vuust & Kringelbach, 2010).

The global prevalence of autism spectrum conditions (ASC, henceforth ‘autism’) is believed to be in excess of 1 in 100 people (Elsabbagh et al., 2012). Autism is therefore a relatively common neurodevelopmental condition, which is characterised by both challenges and strengths. Some of the strengths include huge passion for specific interests that “lie on a continuum with the focused interests of scientists, college professors...” (Jordan & Caldwell-Harris, 2012), great attention to detail (A. Shah & Frith, 1983), and a strong sense of justice (Milner, McIntosh, Colvert, & Happé, 2019), and elevated levels of talent in domains such as art, music, maths and memory (Happé & Frith, 2009). Some of the key challenges for autistic individuals include hyper- or hypo-reactivity to sensory stimuli, such as an aversion to bright lights or loud sounds, or fascination with certain textures or objects, social-communicative and social interaction differences across multiple contexts, and an insistence on and preference for sameness and predictability (APA, 2013).

Understanding the relationship between autism and music listening is a fascinating pursuit for several reasons. Music exposes us to the thoughts, emotions and ideas of others, which can be thought of as a social process. The perceptual mechanisms involved in musical cognition, such as perceiving global and local patterns in the musical contours, are of interest when we consider autistic information processing styles. In one survey of the general population, a majority of respondents reported that their investment in listening to music was principally due to the ability of music to convey emotions (Allen et al., 2009a, 2009b; Juslin & Sloboda, 2001). In autistic listeners, it is therefore of interest to understand how the emotional content of music is processed, especially in the approximately 50% of autistic individuals experiencing alexithymia, or difficulty in perceiving emotions in themselves or others (Berthoz & Hill, 2005; Hill, Berthoz, & Frith, 2004), although it has been suggested that alexithymia is an independent construct with its own neurocognitive and etiological basis (P. Shah, Hall, Catmur, & Bird, 2016).

Several authors have conjectured that individuals with autism may not be emotionally responsive to music (Huron, 2001; Levitin, 2006; Peretz, 2001). The reasoning behind this supposition is that music may have initially evolved for the purpose of promoting social bonding, and that therefore individuals with autism may find it inherently less rewarding than their non-autistic peers without social deficits (Huron, 2001). However, this supposition is not without its critics (P. Heaton, Hermelin, & Pring, 1999). Outstanding musical talent and skill has frequently been reported in autism, including in Leo Kanner’s initial observations of autistic children (Meilleur, Jelenic, & Mottron, 2015). Happé (2018) proposes that detail-focused processing, a notable information processing variance in autism, may underlie autistic musical talent. As Happé highlights, autistic children even without musical training or proficiency are better able to hold exact pitch information in mind for days to weeks, compared to non-autistic children (Pamela Heaton, 2009). In terms of how adults with autism use music in their everyday lives, one semi-structured questionnaire study reported that autistic adults’ uses of music were incredibly similar to that of non-autistic individuals (Allen et al., 2009b). Uses included self-management for emotional states such as depression, to induce mood changes, and for social affiliation.

Processing of any pleasant music in the brain engages limbic and paralimbic regions, as well as regions related to the processing of reward (for reviews see (Koelsch, 2010; Zald & Zatorre, 2011). A few studies have explored musical familiarity using neuroimaging. One PET study used well-known music such as nursery rhymes to test the neural responses of non-musicians (Satoh, Takeda, Nagata, Shimosegawa, & Kuzuhara, 2006). When judging musical familiarity, activation was found in bilateral anterior portions of the temporal lobes, superior temporal regions, and parahippocampal gyri. A later study used fMRI to explore the processing of familiar music compared to unfamiliar music and random tones, with a focus on exploring the “musical lexicon” or the specific memory system that contains all the representations of the specific musical phrases to which one has been exposed, finding focal activation in the right superior temporal sulcus (Peretz et al., 2009). Plailly et al., (2007) compared musical familiarity to odour familiarity, with the goal of exploring whether there is a multimodal neural base of familiarity. For familiar feelings, across music and odour activation pattern was observed in the left hemisphere: the superior and inferior frontal gyri, precuneus, angular gyrus, parahippocampal gyrus, and hippocampus. Using pop and rock music, Pereira et al., (2011) used fMRI to explore how musical familiarity modulates the activity of the brain. They found familiarity to be linked with increased BOLD response in the thalamus, anterior cingulate cortex, nucleus accumbens, putamen and amygdala (Pereira et al., 2011).

A few studies have explored neural processing of music in autism, however, these studies have primarily focused on emotional processing. While listening to happy music, participants with autism have shown decreased brain activity in the premotor area and left anterior insular, compared to TD participants (Caria, Venuti, & de Falco, 2011). Another study using fMRI found that although autistic and TD listeners engaged similar networks when listening to emotional music compared to neutral music, an increased activity was found only for the autistic group when listening to happy music in dorsolateral prefrontal regions and in the insula/rolandic operculum (Gebauer, Skewes, Westphael, Heaton, & Vuust, 2014). The authors suggest that this may reflect increased cognitive processing and physiological arousal in response to emotional stimuli in autism.

Familiar music is unique in that the brain’s predictive mechanisms know what to expect, both in terms of the perceptual properties of the music (“when”) and the emotional content of the piece (“what”). In a predictive coding framework (Friston et al., 2002), the individuals’ priors or topdown predictions for the sensory input will be largely accurate, and there will likely be little prediction error (although no two experiences are the same and prediction error may never be completely eliminated, particularly for complex stimuli). Music is known to be pleasurable when one’s expectations are fulfilled (Gebauer, Kringelbach, & Vuust, 2012), sometimes known as “sweet anticipation” (Huron, 2006; Koelsch, Vuust, & Friston, 2019; Rohrmeier & Koelsch, 2012).

Prediction coding in autism has not been vastly studied experimentally, but has been widely theorised about. Pellicano and Burr (2012) propose that people with autism see the world more accurately – as it really is – as a consequence of being less biased by their prior experiences. The idea that they present is that prior beliefs, which generate top-down predictions, are somehow attenuated in autism, leading to an increased reliance on bottom-up sensory evidence. This would therefore suggest that to the autistic brain, incoming sensory input would seem more novel, and less familiar. In keeping with this hypothesis, autistic listeners have been found to show less auditory adaptation to repeated sounds (Millin et al., 2018), which is perhaps suggestive of an impairment in predictive mechanisms (Sinha et al., 2014). Similarly, Skewes and Gebauer found suboptimal integration of sensory evidence and prior perceptual knowledge in an auditory localization task in adults with autism (Skewes & Gebauer, 2016). Sinha and colleagues (2014) further propose that the autistic preference for “sameness” may arise from the anxiety caused by a world in which unpredictability is elevated. They quote from Dora Raymaker, Director of the Academic Autistic Spectrum Partnership in Research and Education: “insistence on sameness is actually a way of adapting to a confusing and chaotic environment…”

It should be noted that the specific nature of predictive processing in autism has been questioned and is under debate (Palmer, Lawson, & Hohwy, 2017; Van de Cruys et al., 2014). For instance, Friston, Lawson, and Frith (2013) make the suggestion that in autism, there is not a failure of *prediction* specifically, but a failure to instantiate top-down predictions during perceptual synthesis because their assumed *precision* (confidence or reliability of the priors) is too low. The authors consequently label this as a failure of metacognition instead of prediction, which is to say that the autistic brain may not accurately predict the precision of predictions or prior evidence (beliefs about beliefs).

Herein, we investigated the brain’s dynamical repertoire within a small sample of autistic adults compared to TD adults, as they listened to self-selected familiar musical excerpts and a modified (scrambled) version of the same familiar music. We employed a data-driven approach to study the brain dynamics associated with music listening in the two conditions, between groups, focusing upon the dominant functional connectivity (FC) pattern captured by the leading eigenvector of dynamic FC matrices. This method has been successful in discovering key dynamic network differences both at rest (Cabral et al., 2017; Figueroa et al., 2019; Lord et al., 2019) and in task-based data (Alves Da Mota et al., 2020; E.A. Stark et al., 2020). Given the lack of previous findings associated with music familiarity in autism, but knowing the differences associated in emotional, perceptual, sensory, and specifically predictive processing, between autistic and TD individuals, we expected to find between-group differences in emotion, familiarity-related, and reward-related regions.

## Methods

The data analysed in this study is a subsample of a study performed by Gebauer and colleagues (Gebauer et al., 2014).

### Participants

This study included a total of 24 participants, of which 12 had a formal diagnosis of autism. All participants were right-handed and native Danish speakers, with normal hearing. Groups were matched on gender, age, and IQ (Table 1). Participants were IQ-tested using Wechsler’s Adult Intelligence Scale (WAIS-III). All participants with autism were invited for the ADOS testing after the brain scanning session, but unfortunately five participants were unable to come back. However, all participants with autism were previously diagnosed by specialist clinicians and we were given access to their medical records to confirm diagnoses. Participants with autism were medication free at the time of the study and none had comorbid psychiatric disorders. TD participants had no history of neurological or psychiatric disorders. All participants gave written consent. The study was approved by local ethics committee *(De Videnskabsetiske Komitéer for Region Midtjylland)* and it was in accordance with the Helsinki declaration.

**Table 1.** Sample characteristics. Abbreviations: SD – standard deviation; TD – typically developed; F – female; M – male; IQ – WAIS-III full-scale IQ;

|  |  | Autism | TD |
| --- | --- | --- | --- |
| Age | mean | 24.9 | 24.9 |
|  | SD | 5.7 | 5.4 |
| Gender |  | 1F / 11M | 1F / 11M |
| IQ | mean | 104.8 | 112.4 |
|  | SD | 10.1 | 11.6 |

### Paradigm

The paradigm was designed to investigate brain responses to pleasant self-selected (familiar) music. Participants identified their favourite piece of music and heard the 30 seconds that they found most pleasurable, while lying in the scanner. All participants heard the original version of their preferred music first, followed by a scrambled version of the same 30 seconds. The scrambled version was the self-selected music piece played backwards and cut up into small sections that were placed in a random order. Both musical pieces (familiar and scrambled) were repeated five times, and faded for the final 2 sec. Participants were instructed to lie still in the scanner and enjoy the music without doing anything.

### Image acquisition and processing

This dataset was acquired together with data already published by Gebauer and colleagues (Gebauer et al., 2014). All participants underwent the same imaging protocol using a 12-channel head coil in a Siemens 3T Trim Trio magnetic resonance scanner located at the Centre of Functionally Integrative Neuroscience at Aarhus University Hospital, Denmark. An EPI-sequence was acquired with a total of 103 volumes and with the following parameters: TR = 3000 ms, TE = 27 ms, flip angle = 90, voxel size = 2.00 x 2.00 x 2.00 mm, #voxels = 96 x 96 x 55, slice thickness 2 mm and no gaps. An anatomical T1-weighted sequence was performed after the functional scans with the following parameters: TR = 1900 ms, TE = 2.52, flip angle = 9, voxel size = 0.98 x 0.98 x 1 mm, #voxels = 256 x 256 x 176, slice thickness 1 mm, no gaps and 176 slices. Participants used MR-compatible headphones and were instructed to lie still inside of the scanner.

### fMRI Processing

The fMRI data was processed using MELODIC (Multivariate Exploratory Linear Decomposition into Independent Components) (C.F. Beckmann & Smith, 2004) part of FSL (FMRIB’s Software Library, www.fmri.ox.ac.uk/fsl). The first 3 volumes were discarded. The default parameters of this imaging pre-processing pipeline were used for all the 24 participants: motion correction using MCFLIRT (Jenkinson, Bannister, Brady, & Smith, 2002); non-brain removal using BET (Smith, 2002); spatial smoothing using a Gaussian kernel of FWHM 5 mm; grand-mean intensity normalization of the entire 4D dataset by a single multiplicative factor and high pass temporal filtering (Gaussian-weighted least-squares straight line fitting with sigma = 50 seconds). FSL tools were used to extract and average the time courses from all voxels within each cluster in the AAL-90 atlas (Tzourio-Mazoyer et al., 2002).

### Dynamic Functional Connectivity Analysis

We applied a novel method to capture patterns of functional connectivity from fMRI data at single TR resolution with reduced dimensionality, the Leading Eigenvector Dynamics Analysis (LEiDA) (Alves Da Mota et al., 2020; Cabral et al., 2017; Figueroa et al., 2019; Lord et al., 2019; E.A. Stark et al., 2020). On a first stage, the BOLD signals in the *N*=90 brain areas were band-pass filtered between 0.02 Hz and 0.1 Hz and subsequently the phase of the filtered BOLD signals was estimated using the Hilbert transform (Cabral et al., 2017; Glerean, Salmi, Lahnakoski, Jääskeläinen, & Sams, 2012). The Hilbert transform expresses a given signal x as x(t) = A(t)*cos(θ(t)), where A is the time-varying amplitude and θ is the time-varying phase. Given the BOLD phases, we computed a dynamic BOLD phase coherence matrix (dFC, with size NxNxT), where each entry dFC(n,p,t) captures the degree of synchronization between areas n and p at time t, given by the following equation:

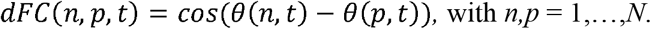

To characterize the evolution of the dFC matrix over time with reduced dimensionality, we considered only its leading eigenvector, *V_1_*(*t*), which is a *Nx1* vector that captures, at time *t*, the projection of the BOLD phase in each brain area into the main *orientation* of BOLD phases over all areas. When all elements of *V_1_*(*t*) have the same sign, all BOLD phases project in the same direction with respect to the orientation determined by *V_1_*(*t*). If instead the first eigenvector *V_1_*(*t*) has elements of different signs (i.e., positive and negative), the BOLD signals project into different directions with respect to the leading eigenvector, which naturally divides the brain into distinct modes (colored in red and blue in Figure 1B middle panels). Previous studies using LEiDA have shown that the subset of brain areas whose BOLD signals appear temporally phase-shifted from the main BOLD signal orientation reveal meaningful functional brain networks (Alves Da Mota et al., 2020; Cabral et al., 2017; Figueroa et al., 2019; Lord et al., 2019; E.A. Stark et al., 2020).

**Figure 1.**
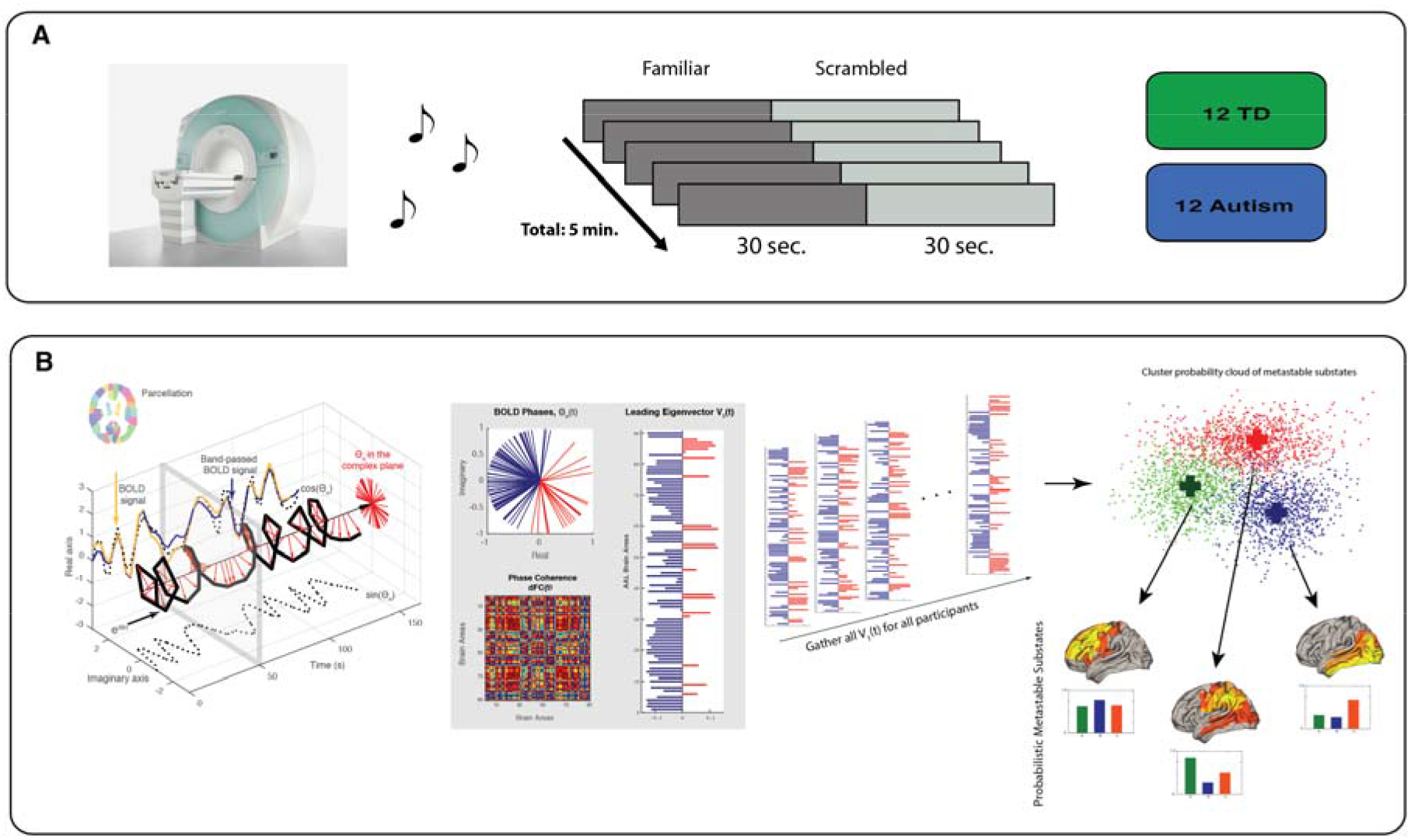
The dynamics of listening to pleasant music (familiar or scrambled), experimental protocol and methods. **A)** Experimental design, participants were asked to listen to a self-selected (familiar) and a scrambled version of the same musical piece whilst inside the MRI scanner. B) Our research question was to understand if individuals with and without autism process familiar music differently, and which brain networks could be driving any differences. We use a leading eigenvector dynamics analysis (LEiDA) to capture the patterns of functional connectivity from fMRI data at single TR resolution (in this study: 3 seconds).

### Stimuli

In order to inform about potential group differences in the self-chosen musical excerpts, we performed a post-hoc exploratory analysis of the acoustical features of all excerpts using the Music Information Retrieval (MIR)-Toolbox in Matlab (Lartillot & Toiviainen, 2007). Due to missing data, this analysis was limited to 10 excerpts from the TD group and 9 excerpts from the autism group. We focussed on dynamic metrical strength, a measure indicating metric predictability that can be computed on the frame-decomposed sound files, to account for the dynamical nature of our fMRI analysis. This measure is defined as the sum of the autocorrelation scores of previously identified relevant metrical levels, i.e. the metrical hierarchy underlying the piece. On a perceptual level, this measure can be thought to represent how clear the pulsation or how strong the metrical hierarchy is, where a large value would reflect a clear, easily recognisable metrical structure, while a smaller value would indicate a more complex or hidden meter (Lartillot et al., 2013). In each group, one outlier was identified in this measure (z-score > 3) and therefore excluded from the group comparison, resulting in 9 and 8 excerpts for the TD and Autism group, respectively. Unpaired two-tailed t-tests between the groups revealed a significantly higher dynamic metrical strength for the autism group (Autism Mean = 0.7067 ± 0.2738 SD; TD Mean = 0.6286 ± 0.1972 SD, p < .0001). However, other (summary) measures that could potentially reflect predictability on other acoustic levels, such as pulse clarity or key clarity, showed no difference between the groups. Additional information on the musical feature analysis can be found in the supplementary material (Figure S1).

### Identification of metastable substates

In this study, we aimed to investigate the existence of specific patterns of functional connectivity, or metastable substates, associated with listening to pleasant familiar music. To do so, we searched for recurrent connectivity patterns emerging between and within groups for familiar and scrambled music and compared their probabilities of occurrence. Recurrent connectivity patterns, or substates, were detected by applying a k-means clustering algorithm to the set of leading eigenvectors, *V_1_*(*t*), associated to the fMRI volumes acquired for all participants. The k-means algorithm clusters the data into an optimal set of *k* clusters, where each cluster can be interpreted as a recurrent substate (Figure 1B right panel).

While resting-state fMRI studies have revealed the existence of a reduced set of approximately 5 to 10 functional networks that recurrently and consistently emerge during rest across participants and recording sites (Christian F Beckmann, DeLuca, Devlin, & Smith, 2005; Cabral et al., 2017; Damoiseaux et al., 2006; Thomas Yeo et al., 2011), the number of substates emerging in brain activity during a task is undetermined, and depends on the level of precision allowed by the spatial and temporal scales of the recordings. In the current study, we did not aim to determine the optimal number of recurrent substates detected in a given condition, but instead to detect substates whose probability of occurrence was significantly different between groups. In that direction, we ran the k-means algorithm varying k from 3 to 15 and, for each k, statistically compared the occurrence of the resulting substates between groups for familiar and scrambled music.

### Probability of occurrence and switching profiles

Recurrent substates were compared in terms of their probabilities of occurrence and switching profiles between groups. Probability of occurrence, which is the number of epochs assigned to a given substate divided by the total number of epochs in each scanning session, was calculated for each substate and for each scanning session. Switching profiles, i.e., probabilities of switching from a given substate to any of the other substates, were also calculated. Statistical differences between groups, in probabilities of occurrence and switching profiles were assessed using a permutation-based unpaired t-test. The significant thresholds were corrected to account for multiple comparisons as *0.05/k,* where *k* is the number of substates (or independent hypotheses) tested in each partition model (Alves Da Mota et al., 2020; Cabral et al., 2017; Figueroa et al., 2019; Lord et al., 2019; E.A. Stark et al., 2020).

## Results

Previous studies have used static functional connectivity, which uses a single measure to quantify the correlation between the BOLD signal of two brain regions over the whole recording time. Instead, in this study, we investigated the dynamic nature of the brain while listening to music, by characterising the ensemble of metastable substates (i.e. the most recurrent patterns of whole-brain functional connectivity) arising during the 5 minutes of exposure to auditory stimuli.

### Detection of significant different substates

Our dynamical analysis revealed seven recurrent substates (k = 7) while listening to pleasant familiar music. A group comparison analysis for k = 7 revealed a recurrent substate with significantly higher probability of occurrence in the TD group than in the autism group (Figure 2). This substate comprises a network with limbic and paralimbic areas, including left: hippocampus and middle temporal gyrus, and bilateral: parahippocampal gyrus and amygdala (Figure 3). These areas are known to play a key role in the processing of emotional music (Koelsch, 2010; Peretz, Aubé, & Armony, 2013).

**Figure 2.**
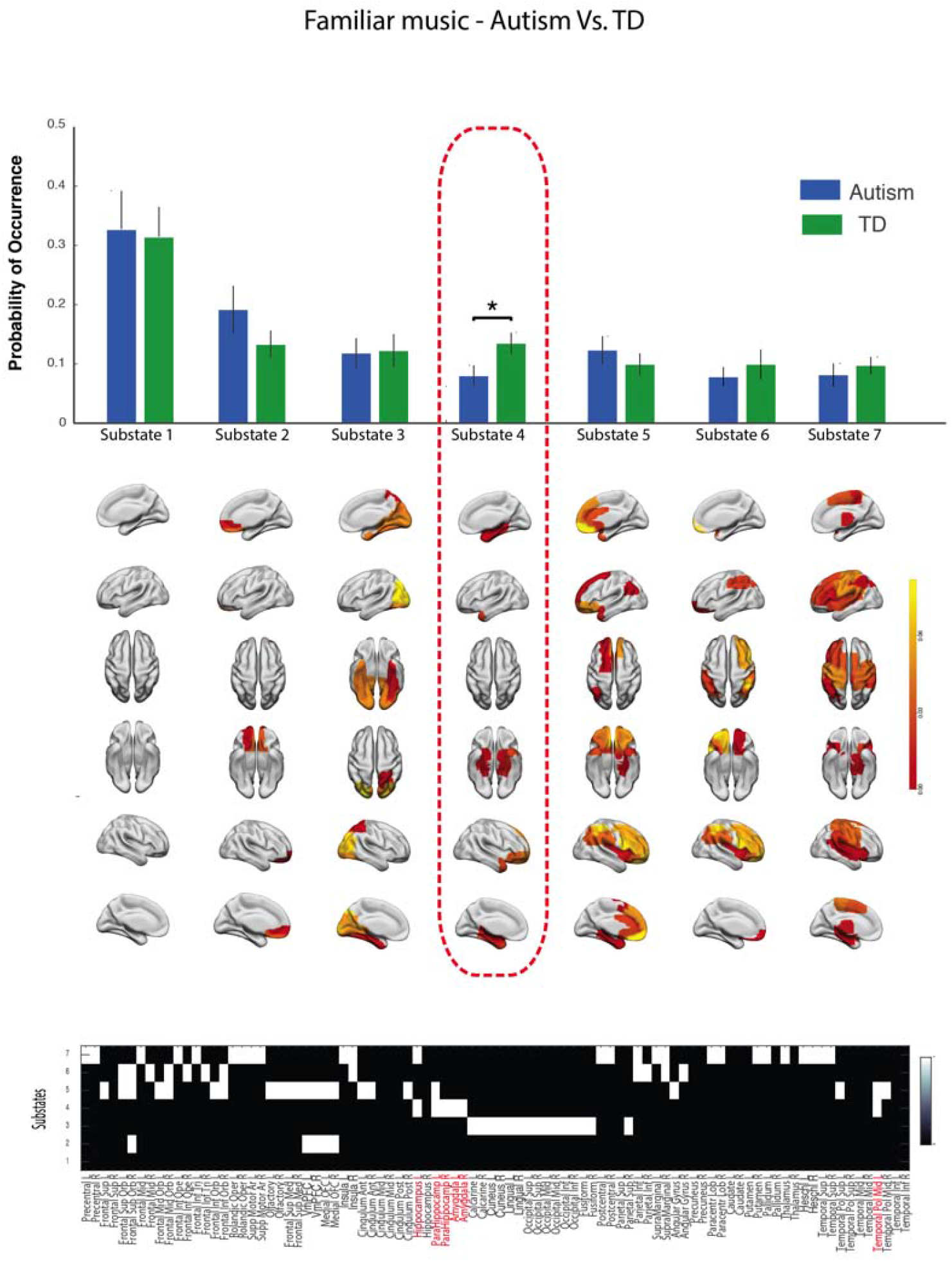
**Repertoire of the seven recurrent metastable substates active while listening to familiar music** (for k = 7). Top: Probability of occurrence of each substate for the autism and TD groups and respective brain rendering of the seven brain substates are represented. Brain substate 4 was found to have significantly lower probability of occurrence for individuals with autism than for TD individuals. Bottom: Nodes belonging to each of the brain substates are represented by a binary matrix, nodes belonging to the substate 4 are highlighted in red.

**Figure 3.**
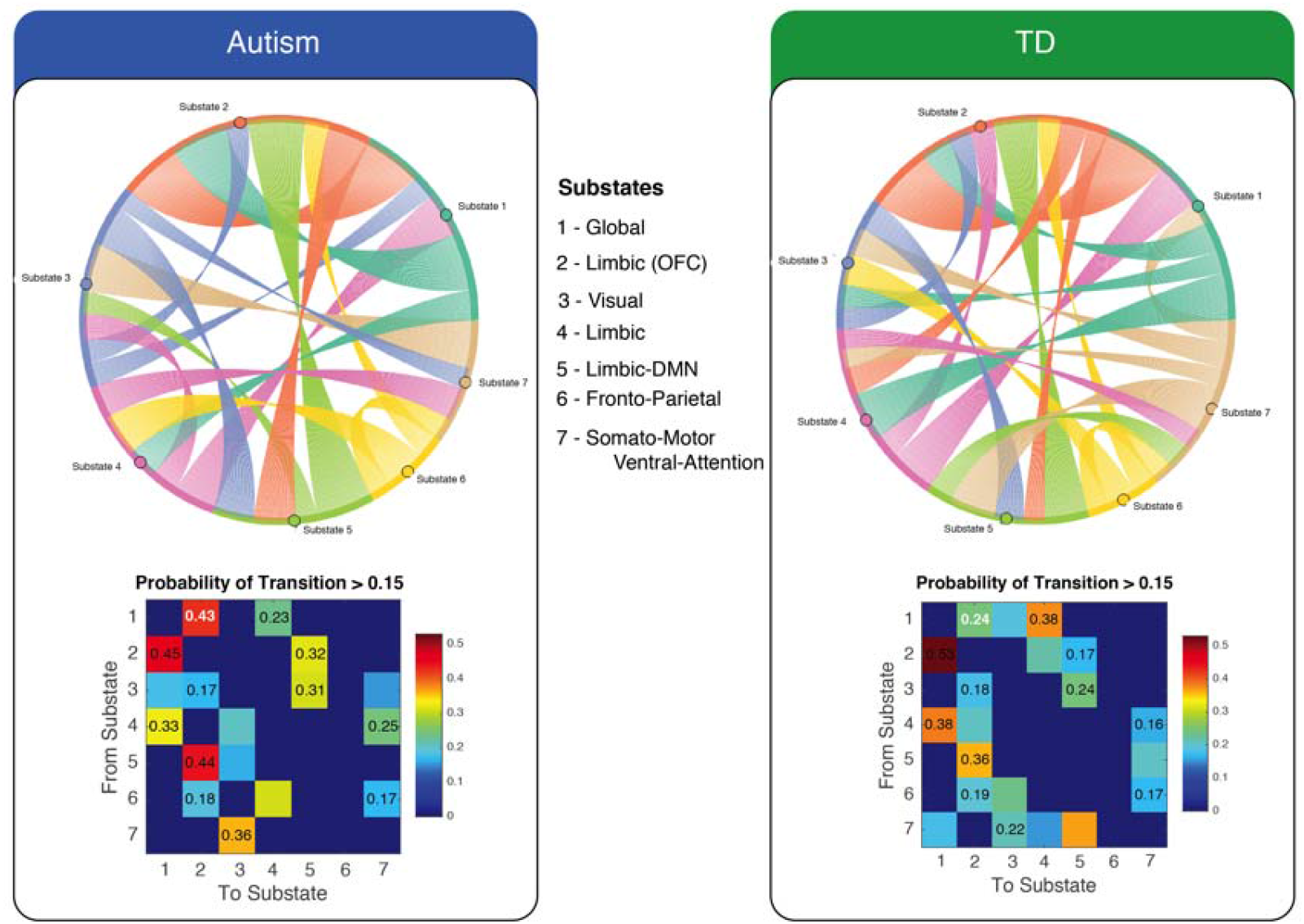
**Individuals with autism have an altered switching profile for listening to pleasant familiar music.** Switching matrices showing the probability of, being in a given substate, transitioning to any of the other substates (left Autism individuals, right TD individuals). Individuals with autism showed a lower probability of switching from the limbic substate 4 to the global substate 1 (and vice versa), and a higher probability to switch from the substate 4 to the substate 7 (SMN-VAN). We found that TD individuals had two exclusive transitions to the limbic substate 4 that did not occur in individuals with autism. No significant differences were found between groups after correction for k = 7. Abbreviations: SMN = somatomotor network; VAN = ventral attention network; TD = typically developing.

No significant differences were found between groups for listening to scrambled (modified) music. Also, no significant differences were found for a within-group comparison for individuals with autism. One substate was found to occur with significantly higher probability for the modified version of the familiar music (scrambled) than for familiar music in TD individuals for a cluster number of 4 (*k* = 4).

### Transition patterns between-groups for familiar music

We further explored the transition patterns between substates by calculating the probability of switching from a given substate to any of the other substates for both groups while listening to familiar music. In Figure 3, switching transitions and probabilities of transition for the autism group and TD group are shown above a threshold of 15% probability of switching to show more frequent transitions. Switching probabilities were calculated for each individual in each group and statistically compared using a permutation-based unpaired t-test (5000 permutations), only if the switch transition occurred in both groups. No significant differences in transition probability were found after correcting for *k*=7 (p< 0.05/7).

We found an altered switching profile in individuals with autism while listening to familiar music, when compared with TD individuals. A lower probability to switch to the substate 4 (limbic network) from the global substate 1 (and vice-versa), and a higher probability to switch from the substate 4 to the substate 7 (SMN-VAN) was observed in the autism group.

## Discussion

The present study investigated the dynamical repertoire of brain networks during listening to pleasant familiar music in individuals with autism and in typically developing individuals. Our study provides a new approach to study how typical and neurodiverse brains process sensory stimuli such as familiar music. To our knowledge, this is the first study investigating the dynamic neural signatures of listening to familiar music in autistic individuals compared to TD individuals. One key aspect of familiar music is the ability of individual listeners to predict the perceptual and emotional content of the subsequent sensory input. We hypothesised that we would find neural differences between autistic and TD listeners due to the fact that individuals with autism may gain great pleasure from “sameness” or familiarity of sensory input, and that predictive mechanisms may differ between groups.

Our study has shown for the first time that the neural dynamics of listening to self-selected familiar music are altered in individuals with autism, compared with TD individuals. We identified a recurrent network, comprising limbic and paralimbic areas, with significantly lower probability of occurrence in individuals with autism. Interestingly, a lower probability of switching to (and from) this particular substate towards a global substate was also found for listening to pleasant familiar music in individuals with autism. The substate with a significantly different probability of occurrence, comprises bilateral parahippocampal gyrus and amygdala, and left hippocampus and temporal pole. These structures are known to be implicated in reward and emotion, and have been identified as key regions of emotional processing of pleasant and unpleasant music stimuli in TD individuals (Koelsch, 2010; Peretz et al., 2013; Salimpoor, Benovoy, Longo, Cooperstock, & Zatorre, 2009). These regions are also known to be activated when listening to familiar music (Pereira et al., 2011).

Understanding what the network does is a tricky endeavour, so we posit some suggestions herein for how the network may act in TD and autistic individuals, focusing upon studies exploring regional functionality and how ablation or damage to each region may affect auditory, emotional, and ultimately musical processing.

The amygdala is a crucial hub involved in many network configurations, and has been implicated in functions ranging from establishing valence or salience, cognition, reward, and social learning (Rutishauser, Mamelak, & Adolphs, 2015). Amygdala activations have previously been found in emotional responses to musical stimuli, namely in listening to familiar music (Freitas et al., 2018; Koelsch, Fritz, v. Cramon, Müller, & Friederici, 2006; Pereira et al., 2011), and in violation of musical expectancies (Gebauer et al., 2012; Koelsch, Fritz, & Schlaug, 2008).

Some research has found that autistic children show hypoactivation in the amygdala in response to social, but also monetary, rewards (Kohls et al., 2012) which the authors suggest is indicative of a general reward processing dysfunction in autism. However, another study reported diminished activation of the amygdala during an emotion matching task in autism, despite intact ability for the autistic participants on the task compared to TD controls (Corbett et al., 2009). This is important to note, as it demonstrates that lower regional activation doesn’t necessarily mean a deficit in processing. In fact, it is more suggestive of differences in processing which ultimately may lead to the same end state (judging the emotions correctly). Unpicking which of the amygdala’s functions may relate to the network herein is tricky, but the clue is perhaps in the connections with other regions.

Activation of the parahippocampal gyrus is described during both emotional arousal (LaBar & Cabeza, 2006) and personal memory recall (Fink et al., 1996) which makes sense given its involvement in the processing of familiar and pleasant music. Unsurprisingly, therefore, the parahippocampal gyrus has subsequently been implicated in processing of music-evoked emotion (Koelsch, 2014), music-evoked personal memories (Janata, 2009), and music mental imagery (Zvyagintsev et al., 2013). Mental musical imagery is the act of imaging music, and is a tool to predict what is likely to happen next in a known song (Herholz, Halpern, & Zatorre, 2012; Zatorre, Halpern, Perry, Meyer, & Evans, 1996), which relies heavily upon predictive processing. Damage to parahippocampal gyrus has been found to lead to patients rating dissonant music, usually rated as highly unpleasant, as “slightly pleasant” (Gosselin et al., 2006), demonstrating its important role in accurate musical perception and valence.

The left hippocampus, present in the key network in this study, has been found to be related to retrieval success for music memory (Gagnepain et al., 2017; Watanabe, Yagishita, & Kikyo, 2008) and to be involved in coding expectancy violations that are caused by reward anticipation (Gebauer et al., 2012; Salimpoor, Zald, Zatorre, Dagher, & McIntosh, 2015). The left hippocampus and temporal pole have been found to be active in familiar music listening contrasted to non-familiar music listening (Freitas et al., 2018; Pereira et al., 2011; Plailly et al., 2007).

The temporal pole lies between the orbitofrontal cortex and the amygdala and receives and sends connections to both regions (Olson, Plotzker, & Ezzyat, 2007). It has been described as an “association cortex” due to its ability to both receive and send connections to the three sensory systems, including auditory information, represented in the temporal lobe. Olson, Plotzker, and Ezzyat (2007) propose that one role of the temporal pole is to “bind complex, highly processed perceptual inputs to visceral emotional responses”. One interesting case study comes from a patient who had a left temporal lobectomy, and subsequently underwent a drastic change in his musical preferences, from rock music to Celtic or Corsican polyphonic singing (Sellal et al., 2003). This case study demonstrates that the temporal pole has a role in musical preferences and emotional processing of complex auditory information, and perhaps its role is facilitated by its network connections to limbic regions such as the amygdala and mnemonic regions such as hippocampus and parahippocampal gyrus.

All regions in the aforementioned network thus appear to be involved in the processing of *familiar* music specifically (Freitas et al., 2018; Pereira et al., 2011), be it associating the sensory information to emotional responses, familiar music recall or establishing salience (a process highly dependent upon memory). The reason why this key network may have lower probability of occurrence in autistic listeners may be due to differences in predictive processing. As Pellicano and Burr (2012) suggest, the autistic brain may have attenuated top-down predictions, making familiar stimuli seem more ‘novel’. This specific network, which is heavily linked to the processing of familiar music, may have a lower probability of occurrence in the autistic listeners simply because they engage less with activating priors from memory, and forming top-down predictions. This is not to say that the music is not still rewarding to the autistic listeners, simply that they may activate fewer priors or be less confident in the precision of their priors (Friston et al., 2013; Pellicano & Burr, 2012).

Literature shows evidence of impairment of predictive abilities in autism (Lawson, Rees, & Friston, 2014), however the specific nature of the possibly attenuated predictions in the autistic group is an interesting and, as yet, unanswerable question. Such top-down predictions may be separated into “what” and “when” predictions of auditory events, or into perceptual properties (i.e. the next chord should be in C major), lexical content if the music has a vocal element, or emotional content (the upcoming key change may induce “chills”).

Rohrmeier and Koelsch (2012) proposed that predictive information processing is fundamental to music in three different ways: (1) dynamics of musical temporality, (2) musical interaction and synchronisation, and (3) in inducing specific emotional and aesthetic musical effects, such as pleasure or “chills.” Given the profile of autistic strength in “local” or detail-focused processing (A. Shah & Frith, 1983) and for sublime feats of memory for extended melodic pieces (Stanutz, Wapnick, & Burack, 2014), as first described by Kanner (Kanner, 1943, 1971), it remains in question as to whether the autistic group may have difficulties in aspect one. The study involved no semblance of interaction or synchronisation, so aspect two may also be unrelated in this case. Aspect three, the induction of specific emotional and aesthetic musical effects, may be where the predictive differences lie, although again we cannot tell from this specific dataset. Despite some suggestions of dysfunctional emotional processing during music listening in autism (Caria et al., 2011), other studies have found intact emotional processing (Gebauer et al., 2014) in terms of assessing the valence of musical excerpts. However, whether, or to what extent, individuals activate emotional priors, or have confidence in the precision of these priors, when listening to music remains an open question. An elegant means of studying this would be to explore brain activity *and* behavioural ratings during conditions where the emotional predictability of a musical piece changes over time, while also measuring alexithymia as a covariate.

Individuals with autism may have difficulties in perceiving the unpredictable world, therefore leading to an “insistence on sameness” and reluctance to experience change or new experiences (APA, 2013). In the context of familiar music, although the autistic listeners may activate fewer top-down priors, the music may be especially pleasing for its relative predictability. Our results also show that the ventromedial prefrontal cortex and medial orbital frontal cortex have a higher probability of occurrence in individuals with autism, when compared with TD individuals (substate 2). These limbic regions may be responsible for online evaluation of the value of music and for attributing reward value (Salimpoor et al., 2015). Their higher probability of occurrence may be evidence of “sweet anticipation” and high reward values within the autistic group.

It has been proposed before that the process of listening to emotional music has a specific temporal dynamic, as music evolves over time (Koelsch et al., 2006). Studying the process of such dynamic stimuli cannot be assumed to be temporally stable over the whole scanning time. Dynamic functional connectivity aims to characterize the complex temporal changes in the network structure while participants listen to the musical excerpts. Here, we relate these temporal changes in the brain to dynamic measures of the musical pieces. Recent studies have explored the dynamic FC (dFC) patterns in resting-state in individuals with autism and have found different results than the ones found in static FC analysis (Chen, Nomi, Uddin, Duan, & Chen, 2017; de Lacy, Doherty, King, Rachakonda, & Calhoun, 2017; Falahpour et al., 2016; Watanabe & Rees, 2017). These studies reported that the autistic brain shows fewer state transitions than the typically developing brain (de Lacy et al., 2017; Watanabe & Rees, 2017) and dFC analysis have been also found to improve diagnostic prediction in autism (Zhu et al., 2016). However, no study has yet explored the dynamic connectivity differences associated to sensory stimulation such as with music.

This work highlights several promising routes for further exploration, but also has several limitations which will be important to mitigate in future work. First, our substates are constrained by the parcellation atlas selected, the AAL 90. Extending this methodology to encompass parcellation schemes based upon a higher number of regional areas may reveal more fine-grained substates (Craddock et al., 2013; Glasser et al., 2016; Shen, Tokoglu, Papademetris, & Constable, 2013). Second, the functional meaning of the substates still remains unclear although we can draw reverse inferences from inputting the brain coordinates into databases such as Neurosynth (Yarkoni, Poldrack, Nichols, Van Essen, & Wager, 2011) or from literature searches where the same brain region has been found to be active under certain conditions. Building whole-brain computational models which can be perturbed in-vivo may offer a promising route towards isolating the role of a given region within a large-scale network (Deco & Kringelbach, 2014).

As has been suggested by MEG studies, meaningful brain activity is thought to occur at time scales of approximately of 200ms (Baker et al., 2014; Vidaurre et al., 2016). Importantly, prediction also occurs on multiple timescales, from milliseconds to minutes, and in multiple hierarchical descriptions (Kiebel, Daunizeau, & Friston, 2008). Since our temporal resolution (TR=3 seconds) did not allow us to capture more rapid dynamics, we limited our analysis to the probability of occurrence and switching profiles occurring over time. Lastly, it would be of great importance to extend this approach to a higher sample size, with different age groups and greater numbers of both genders. A resting state scan of the same population would also allow an interesting comparison to task-based activity.

To conclude, we have discovered preliminary evidence of dynamic network differences in autistic listeners compared to typically developing listeners when hearing familiar, pleasant music. Although the specific role of these networks in familiar music listening remains undetermined, we speculate that it might be due to two elements: primarily predictive processing differences between autistic and TD listeners, but also differences in value attribution due to the frequent autistic need for sameness and predictability. The priorities for future research in this area are to experimentally establish the nature of autistic predictive processing differences more generally, to isolate processes of prediction during music listening into meaningful categories such as temporality, interaction and synchronisation, and emotional predictions (Rohrmeier & Koelsch, 2012), and to resolve dynamic network differences with finer temporal specificity.

